# Meta-analysis of RNA-seq gene fusion detection tools: performance and variability across benchmarks

**DOI:** 10.1101/2025.01.20.633905

**Authors:** Iga Ostrowska, Tomasz Gambin

## Abstract

In this study, we conducted a comprehensive meta-analysis of ten publicly available benchmark tests evaluating the performance of gene fusion detection tools using RNA-seq data. Our analysis focused on key performance metrics, including sensitivity, precision and F1 scores. We evaluated how the tools perform in different data sets. We examined the impact of data set characteristics such as sample type (real or simulated) and read length, as well as additional sample and sequencing parameters on the results. In addition to evaluating performance, we analyzed the organization and design of the benchmark tests, highlighting good practices such as clear descriptions of the datasets, detailed instrument parameters and transparent methodologies. However, we also identified common pitfalls, including insufficient reproducibility information, limited diversity of datasets, and the lack of widely accepted gold standard datasets. These limitations make it difficult to consistently evaluate tools and compare across benchmarks. By synthesizing these findings, we make recommendations for future benchmark projects, emphasizing the need for standardization, increased transparency and the development of robust truth sets. This study aims to help the community create more reliable and reproducible benchmark tests, ultimately accelerating the development and evaluation of gene fusion detection tools for clinical and research applications.

## Introduction

Gene fusions (presented on Figure 1), molecular event which result from chromosomal rearrangements, are a significant focus in molecular biology and genomics, particularly in the study of cancer and genetic diseases. Fusion genes can be caused by various chromosomal aberrations, including translocations, duplications, inversions, or small interstitial deletions. These fusions occur when fragments of different genes combine, leading to the creation of hybrid transcripts. This can result in the production of abnormal proteins that disrupt cellular functions, contributing to processes like uncontrolled cell proliferation and resistance to apoptosis. As a result, many gene fusions have become important targets for targeted therapies, underlining their importance in clinical oncology. Numerous fusion genes have been identified as key drivers in various human cancers. Hematopoietic malignancies, in particular, are well-characterized by the abundance of fusion genes. For instance, in chronic myeloid leukemia, the BCR-ABL1 fusion is present in over 95% of cases, while acute promyelocytic leukemia is characterized by the PML-RARA fusion in 90% of cases. In acute myeloid leukemia (AML), fusion genes are present in about 30% of patients and are often used as critical markers to define clinically relevant subtypes.^1–4^ The discovery and analysis of gene fusions are crucial for advancing precision medicine, facilitating molecular diagnosis, disease prognosis, and the development of novel therapies. Furthermore, gene fusions serve as valuable biomarkers in screening tests and monitoring treatment responses, thereby enhancing the overall quality of patient care. Advancements in RNA sequencing (RNA-seq) technology have made gene fusion detection more accurate and widely accessible. RNA-seq enables comprehensive transcriptome profiling, allowing for the identification of rare and atypical gene fusions that may be overlooked by traditional techniques such as fluorescence in situ hybridization (FISH) or RT-PCR. However, the reliability of RNA-seq results often depends on the algorithms and bioinformatics tools employed in the analysis.^1^

**Figure 1.**
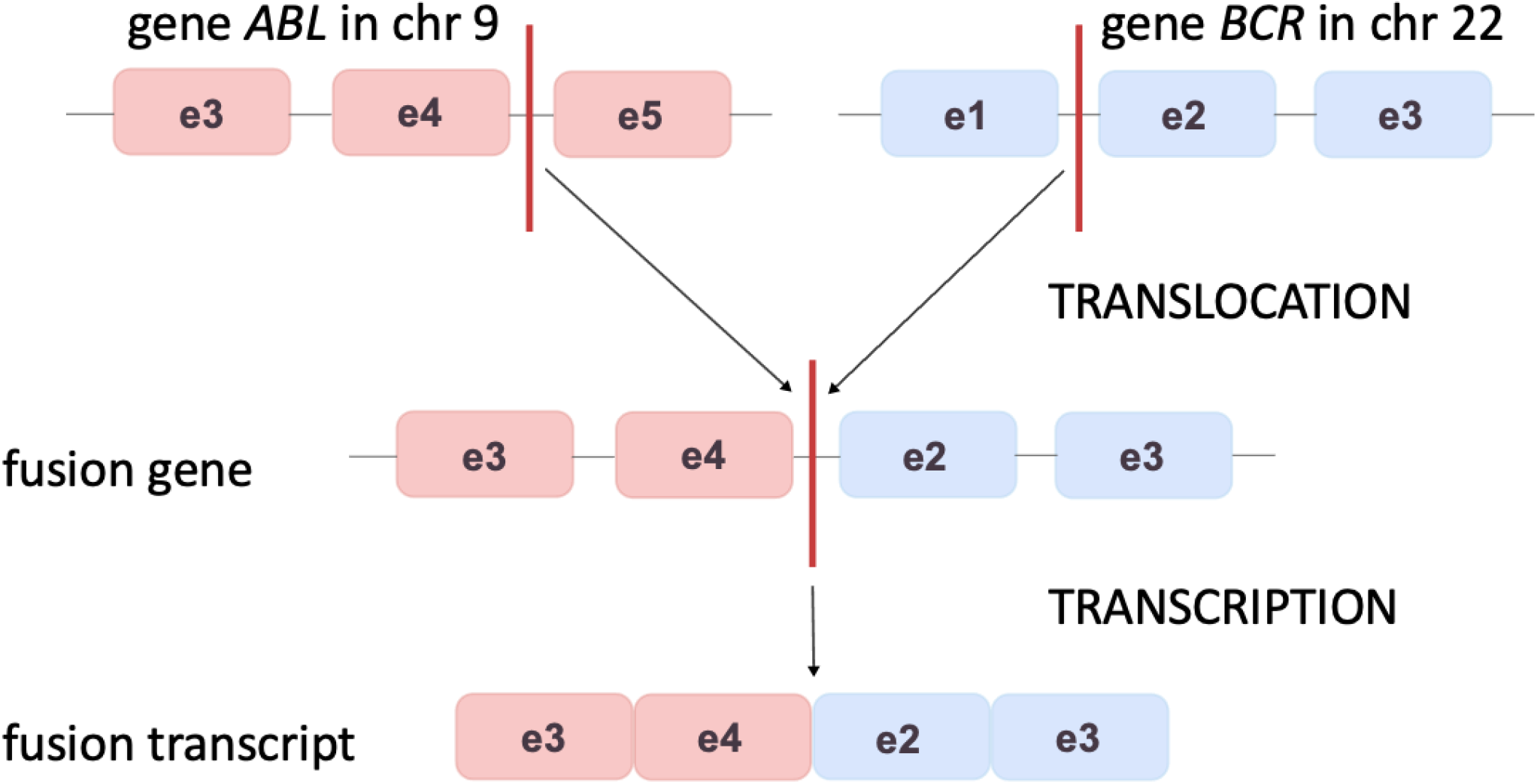
The process of formation of a fusion gene and a fusion transcript as a result of chromosomal translocation between the ABL gene on chromosome 9 and the BCR gene on chromosome 22. The top panel shows the original locations of the exons of both genes in the genome (e3-e5 for ABL, e1-e3 for BCR), with the location of the translocation marked by red lines. This event results in the formation of a fusion gene, in which the e3 and e4 exons of the ABL gene are fused to the e2 and e3 exons of the BCR gene. The bottom panel illustrates the transcription process leading to the formation of the fusion transcript, which contains sequences from both genes. The graphic visualizes the molecular mechanism behind the formation of fusion transcripts, which are key biomarkers in cancer diagnosis and research, such as chronic myeloid leukemia.

There are currently many bioinformatics tools dedicated to the detection of gene fusions. Each of these tools uses different strategies to analyze RNA-seq data, including mapping to a reference genome, identifying inconsistencies in sequence reads, and analyzing evidence supporting the presence of fusions.^5^ The variety of these methods results in significant differences in their sensitivity, precision, and robustness to noise in the data. Tools are often tailored to operate on specific data and cope better with a given data set or technology. Benchmarking plays a critical role in the evaluation of these tools. By providing standardized datasets and objective criteria for comparison, benchmarks enable researchers to assess the performance of gene fusion detection methods in a controlled and reproducible manner. They facilitate the identification of strengths and weaknesses in the algorithms, highlight their applicability to specific biological contexts, and uncover limitations in handling noise or edge cases. Furthermore, benchmarks serve as a foundation for the development of new tools by defining performance expectations and promoting healthy competition within the bioinformatics community.

The goal of this study is to perform a meta-analysis of existing gene fusion detection tools by synthesizing the findings of ten published benchmarks. We aim to evaluate the performance of these tools across different datasets, focusing on key metrics such as sensitivity, precision, and F1 score. We also note the diversity of datasets on which the tools are tested. Additionally, we highlight the strengths and weaknesses in benchmark design and suggest guidelines for future studies.

## Results

### Tool Performance Across Benchmarks

In this paper, we evaluated a diverse set of benchmarks (^5–14^) encompassing 41 tools tested on transcriptome data across 10 benchmarks using 38 datasets (unique or duplicated across benchmarks, detailed in Table 1). These benchmarks represent a broad spectrum of methods and datasets, ensuring a comprehensive evaluation of tool performance. Detailed information on the tools and references is provided in the Supplementary Material. We summarized the number of tools and datasets included in each benchmark, along with additional benchmark features such as the types of datasets used, and the availability of information regarding tool versions, runtime, and memory usage. Detailed information on the tools and datasets tested in each benchmark is provided in Figure 2. Other benchmark features, such as tool versions, dataset types, runtime, and memory usage, are included in the graphic. We observed that the performance of gene fusion detection tools varies significantly across comparative tests using different types of data. Key challenges for the tools being compared include the source of the sample (real or simulated) and the length of the sequencing reads, both of which have a substantial impact on performance metrics. The lack of well-established and comprehensively validated gold standards for fusion detection datasets also influences the results. For this reason, we avoided direct comparisons between results for different cancer types and library preparation methods (e.g., WTS (Whole Transcriptome Sequencing) versus targeted). Additionally, due to the very limited availability of single-end sequencing results (present in only one benchmark), these were excluded from comparisons with paired-end data. It is worth noting that some tools do not support single-end data, further limiting their use in such analyses.

**Table 1.**
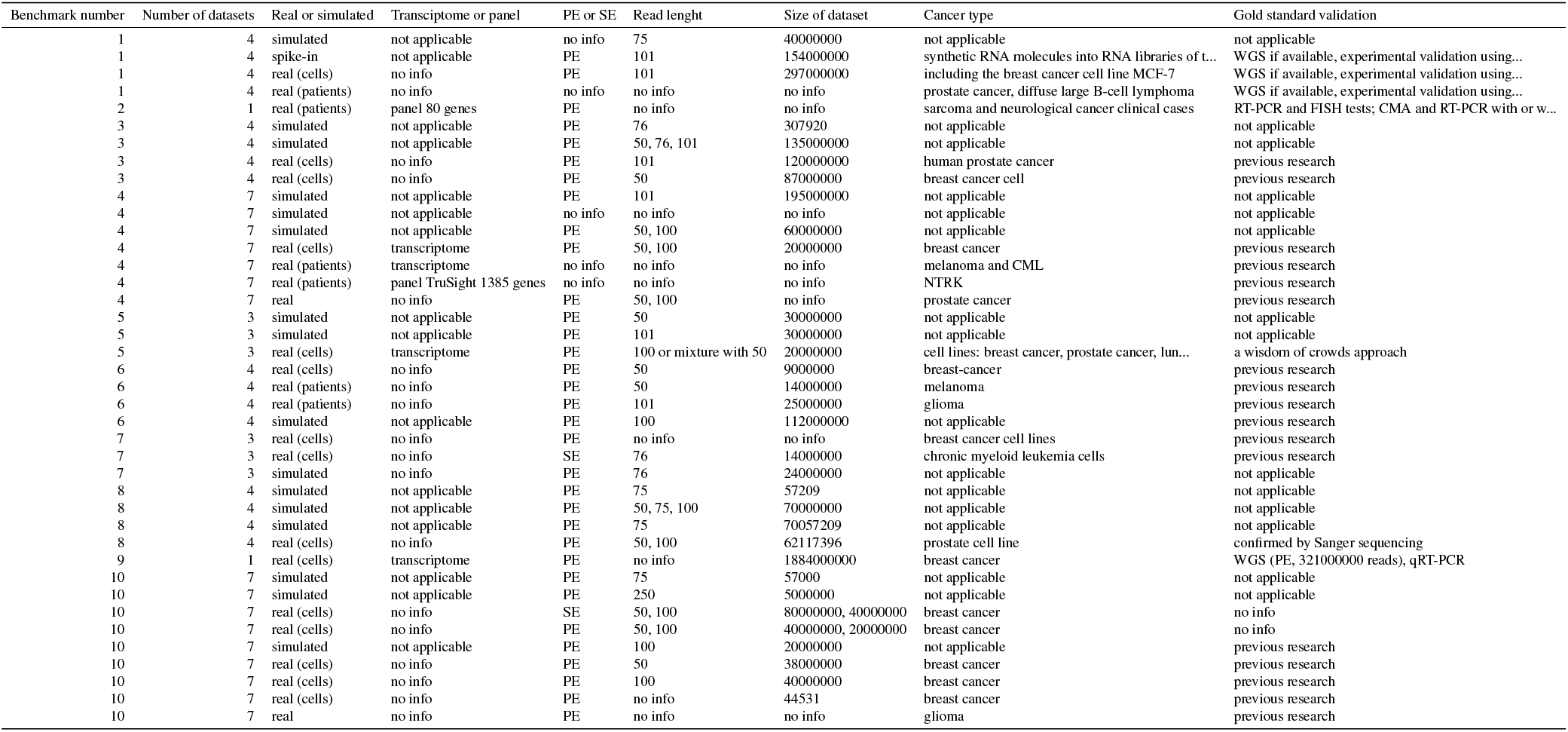
Comparisons of datasets used in benchmarks. The table includes the benchmark number, the number of datasets in a given benchmark, the origin of the sample - real or simulated, the type of library - whole transcriptome sequencing or targeted sequencing (gene panels), the type of sequencing PE - paired-end or SE - single-end, the size of the dataset, the type of cancer whose RNA was sequenced and the type of validation of the truth set, the so-called gold standard.

**Figure 2.**
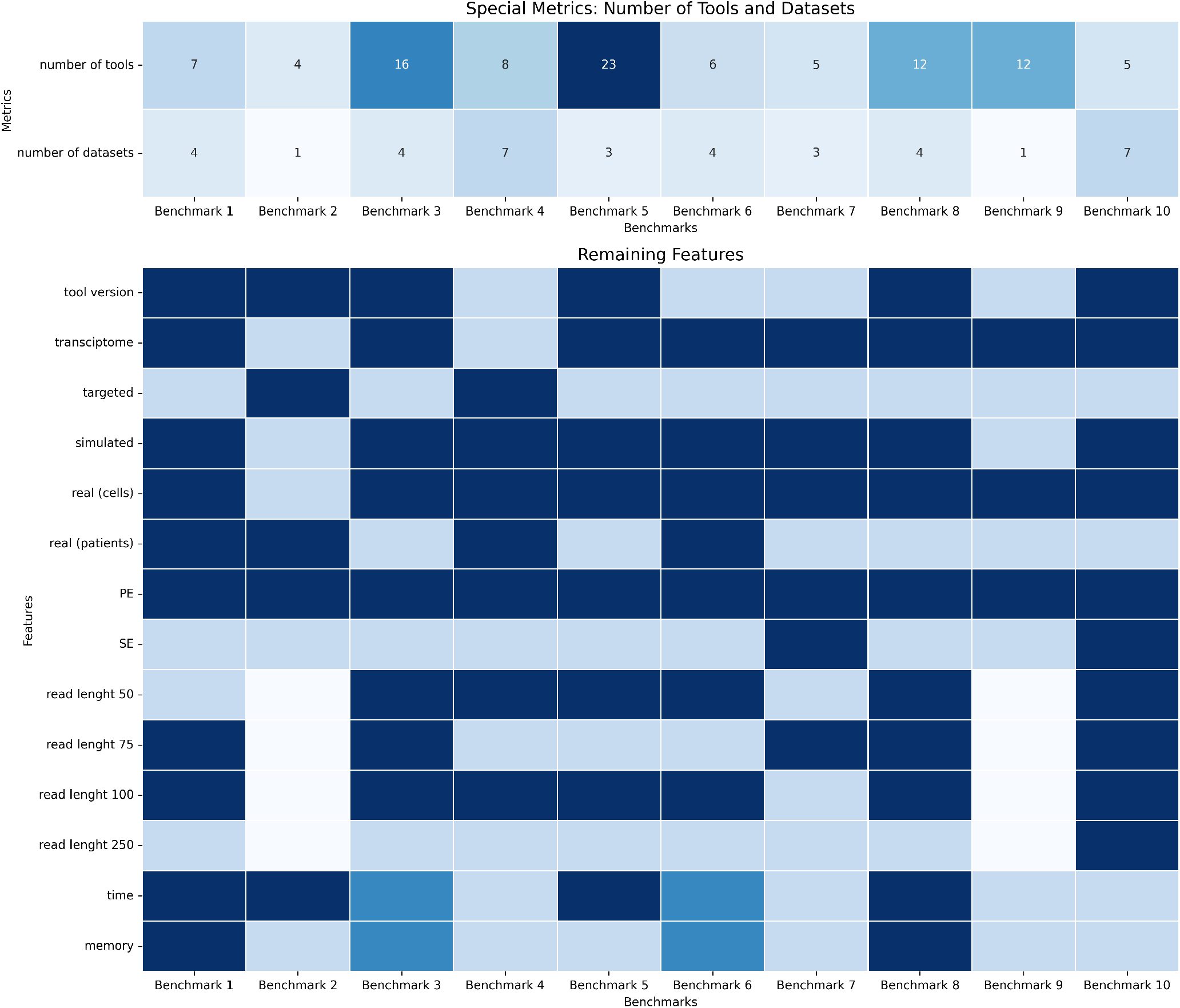
Description of benchmarks included in this study. The Metrics section includes the numbers of tools used and datasets analyzed in benchmarks developed by Uhrig et al.^6^, Balan et al.^7^, Singh et al.^8^, Apostolides et al.^9^, Haas et al.^5^, Vu et al.^10^, Zhao et al.^11^, Zhang et al.^12^, Kumar et al.^13^, and Davidson et al.^14^. The Features section includes the other benchmark parameters. The color gradient of the blue shades depends on the values of the included features (dark blue - feature included, full information; medium blue - feature included, partial information; light blue - feature not included, white - no information about the given feature). “Tool version” refers availability of the specification of tool versions in the benchmark, “Transciptome” and “Targeted” indicate the type of analyzed samples in the datasets, considering full transcriptomic and targeted sequencing. “Simulated”, “real (cells)” and “real (patients)” reflect the origin of the samples in the datasets. “Simulated” are considered to be sets generated by simulators. The real data sets can have two sources: from sequencing from cultured cancer cell lines or from patient samples, derived from actual human RNA sequencing. “PE” and “SE” denote the sequencing type, which are paired-end and single-end sequencing respectively. “Read length” takes into account the sequencing lengths - 50, 75, 100 and 250 bases. The “time” and “memory” features indicate whether the benchmark takes into account the analysis time of the tools, as well as memory usage.

#### Truth set differences

We compared the performance of tools with respect to the type of cancer. The most popular cancer type of real samples tested in the benchmarks is breast cancer (8 out of 10 benchmarks used data from this cancer, see supplements for details). The most popular dataset for breast cancer is the one from the publication by H. Edgren^15^, as it is used by 7 out of 10 benchmarks. For this dataset, the F1 scores differ significantly (see Figure 3), with high standard deviation values (standard deviation from 0.04 for deFuse^16^ to 0.20 for Arriba^6^). We hypothesize that these differences could be explained by multiple factors, such as above mentioned lack of well validated golden standard. In fact, the number of reported true positives for the single dataset varies from 27 to 99 in 7 different benchmarks.

**Figure 3.**
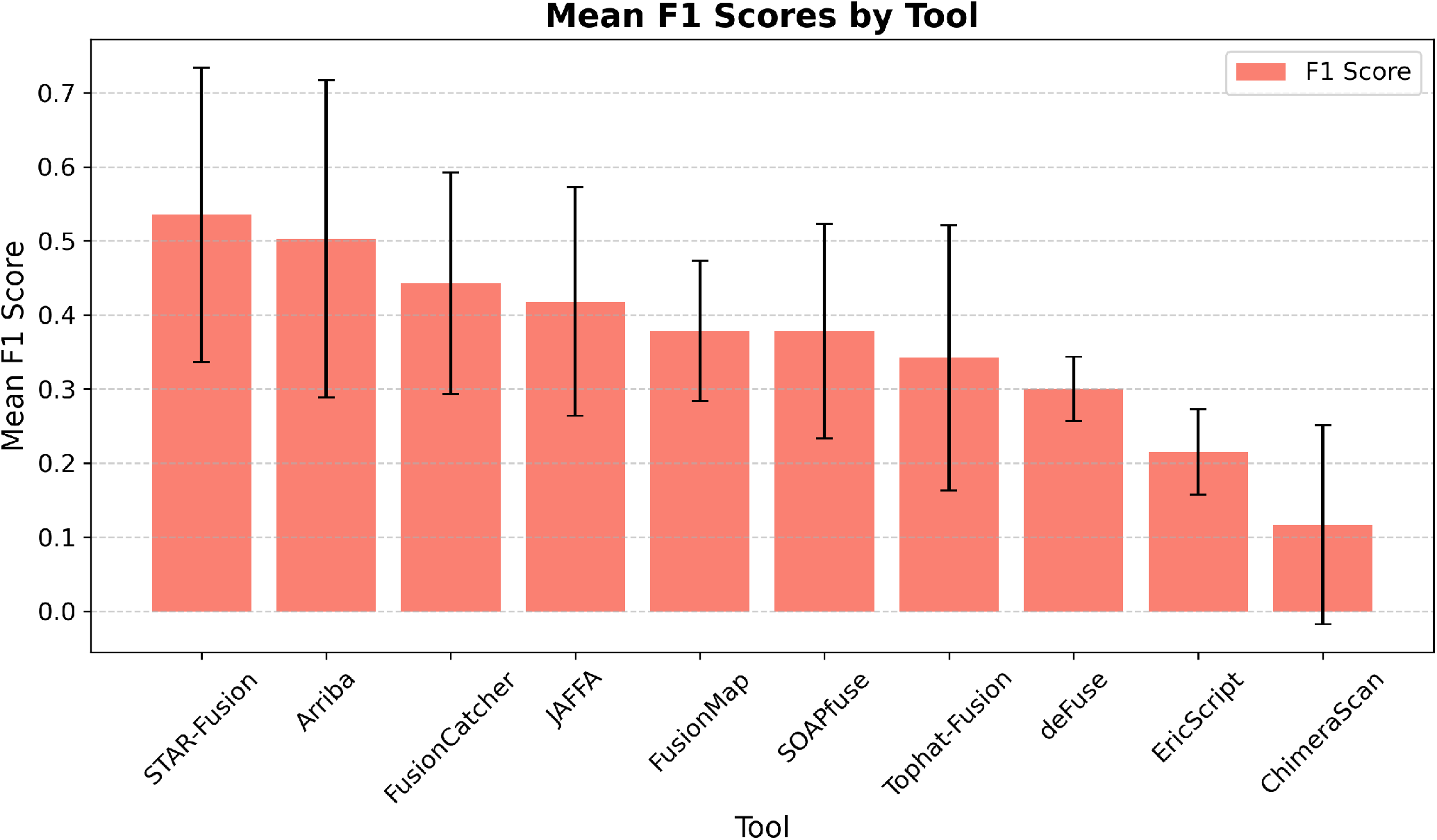
F1 score with standard deviation (error bars) for each gene fusion detection tool on the Edgren dataset.

**Figure 4.**
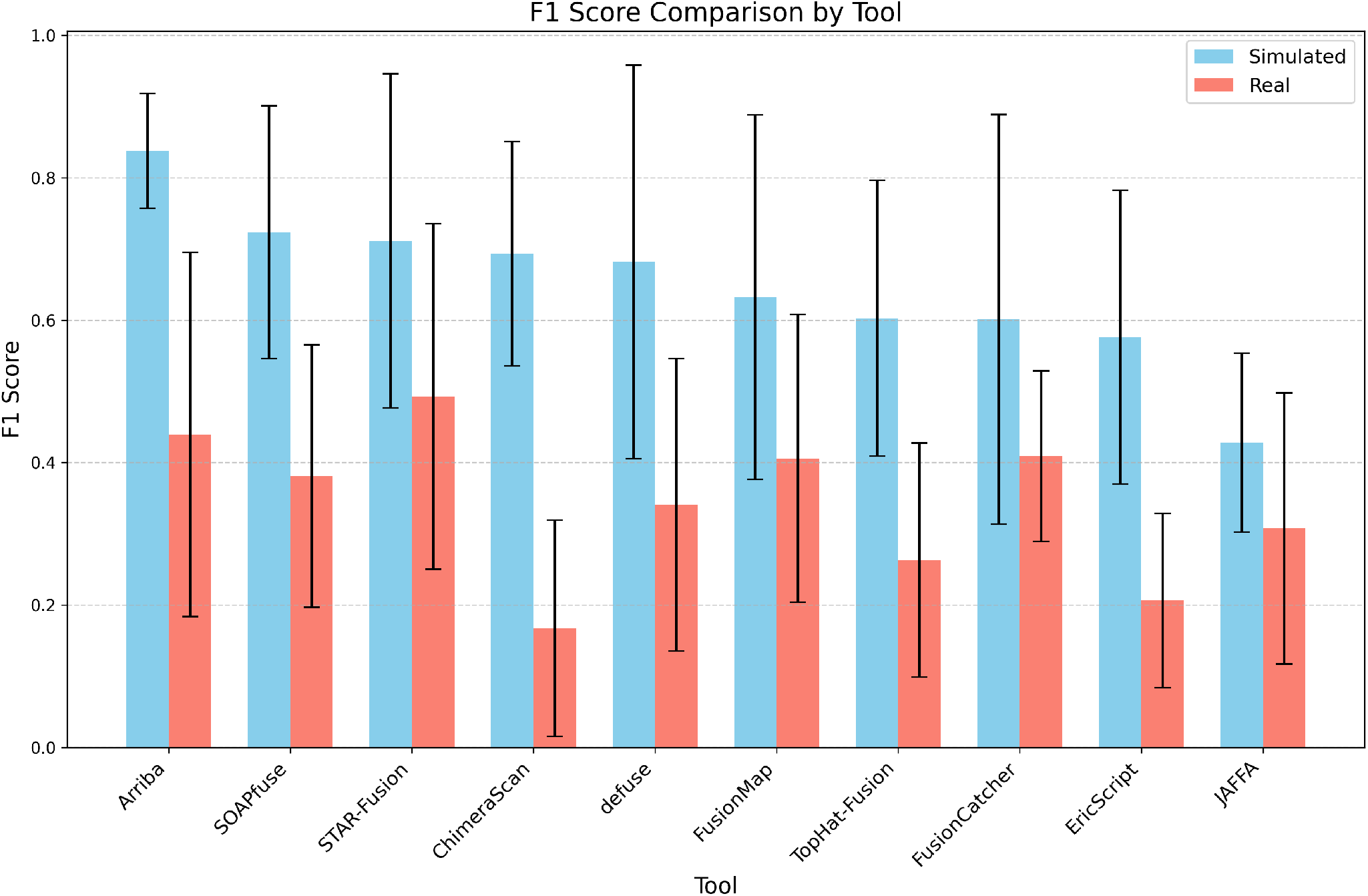
Comparison of F1 score with standard deviation (error bars) for real (red) and simulated (blue) samples for each gene fusion detection tool.

#### Simulated vs. Real Datasets

For the analysis of the influence of the sample source, tools with at least five entries available for both types of datasets were selected. To create the plot, the performance of 10 tools was compared: Arriba^6^, SOAPfuse^17^, STAR-Fusion^5^, ChimeraScan^18^, defuse^16^, FusionMap^19^, TopHat-Fusion^20^, FusionCatcher^21^, EricScript^22^ and JAFFA^14^. Analyses using real datasets in comparison to simulated datasets (shown on the plot 4) resulted in lower F1 scores, likely due to inherent biological complexity and variability that simulated datasets may not reflect. Commonly occurring noise in biological samples definitely causes underestimation of results. Another cause of lower results for real samples may be the previously mentioned problem with validating the truth set, which obviously does not apply to simulated samples.

#### Read Length

To achieve a more reliable comparison, only simulated data were used. Two results for 50 bp and 100 bp lengths were used to create the plot (same two comparison benchmarks for both lengths). Figure 5 compares the following 5 tools: STAR-SEQR^23^, STAR-Fusion, Arriba, deFuse and EricScript,. Shorter reads (50 bp) resulted in lower F1 scores in most tools, indicating the importance of read length in accurate fusion detection. Some tools, such as FusionCatcher, FusionMap, JAFFA, and TopHat-Fusion, did not work with 50 bp reads. This limitation may be specific to certain versions of these tools. Detailed information on tool versions and their compatibility is provided in the Supplementary Materials. Other tools were successfully used in 50 bp benchmarks but did not meet the comparison criteria. For instance, they were tested only on real samples or lacked corresponding 100 bp data in simulated samples for direct comparison.

**Figure 5.**
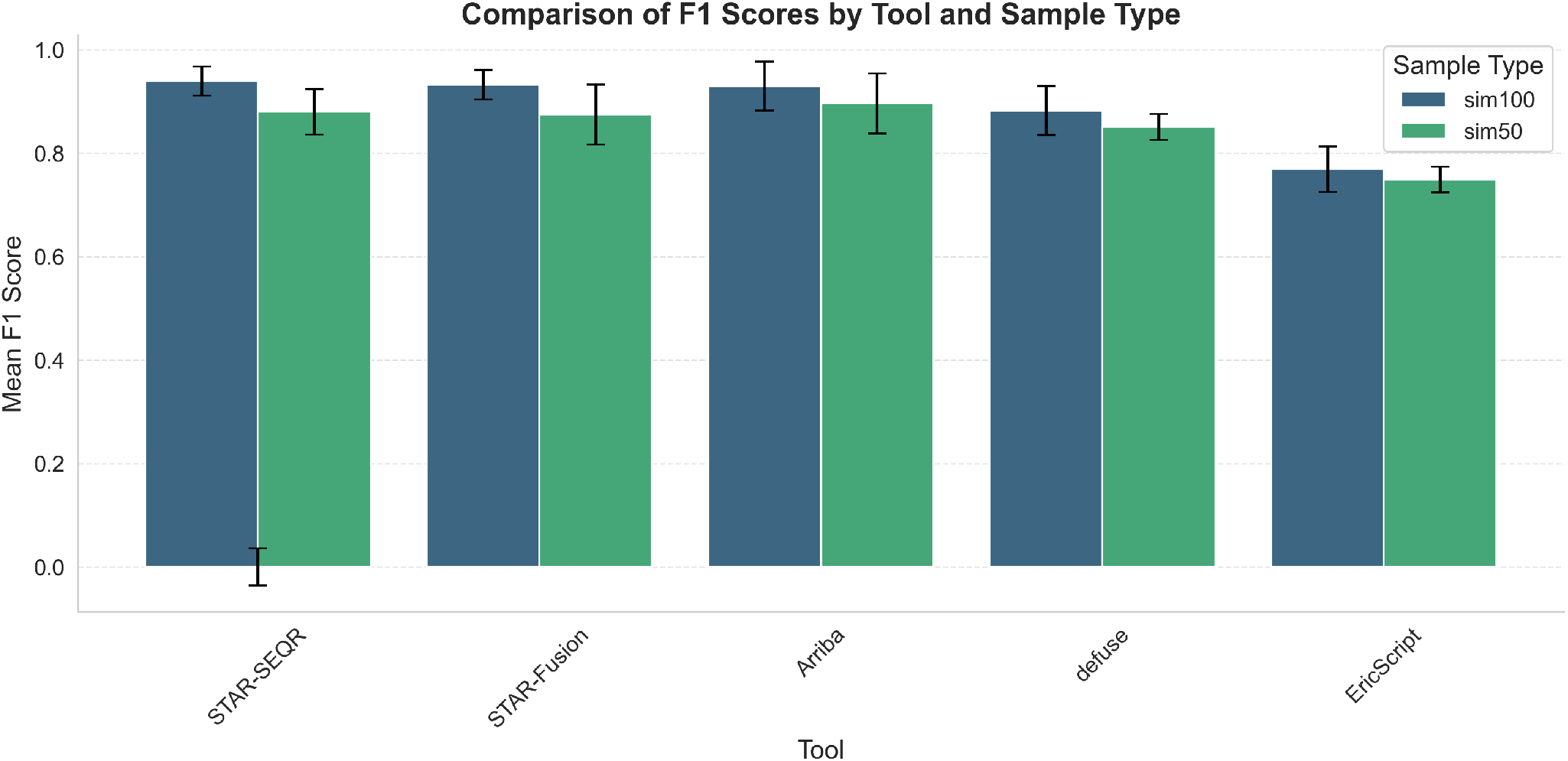
Comparison of F1 score with standard deviation (error bars) for simulated 50bp (green) and simulated 100bp (blue) datasets for each gene fusion detection tool.

### Key observations

The results presented in the previous subsections represent a selection of the findings from this meta-analysis, with additional details available in the Supplementary Data. One significant limitation identified is the incomplete documentation in many studies. As shown in Figure 1, only 50% of benchmarks reported critical elements such as tool versions, parameter settings, and dataset composition. This lack of transparency undermines reproducibility and credibility, as researchers attempting to replicate benchmarks often encounter inconsistencies or missing information.

Another issue is the limited diversity of datasets used in benchmarks. While all benchmarks included real clinical samples, only 8 out of 10 incorporated simulated datasets, which are crucial for controlled performance evaluations. Additionally, there were notable differences in read lengths across benchmarks, with 8 out of 10 relying exclusively on data from whole-transcriptome sequencing (WTS) rather than targeted sequencing approaches. This narrow focus on WTS datasets does not fully capture the variability and challenges presented by targeted sequencing data, potentially skewing tool performance assessments.

A further notable issue is the frequent inclusion of “filler tools” in benchmark studies. These are tools that appear across multiple benchmarks, often without sufficient justification for their inclusion. For example, TopHat-Fusion^20^ was included in 8 out of the 10 studies analyzed, despite known limitations and its outdated status relative to newer methods. While these tools often deliver average performance—neither excelling nor performing poorly—they tend to dominate benchmarking comparisons. Their frequent inclusion creates a skewed representation of tool performance and does not necessarily advance the identification of the most effective or innovative methods. This overrepresentation of filler tools often comes at the expense of including more recently developed or specialized tools that might offer improved performance but are less commonly used. It also limits the diversity of approaches being evaluated, potentially slowing the adoption of newer, more effective solutions. Addressing this imbalance requires a more deliberate selection of tools in benchmark studies to ensure fair and comprehensive evaluations. Additionally, computational metrics such as runtime, memory usage, and scalability are often overlooked in benchmarks, despite their importance for clinical applications where computational efficiency is crucial. Moreover, the methodologies, evaluation metrics, and dataset preparation approaches used in different benchmarks vary widely, making it challenging to compare studies. This inconsistency underscores the need for standardized benchmarking practices to enhance reliability and comparability across studies.

## Discussion

The meta-analysis revealed significant variability in the design, execution, and reporting of benchmarks for gene fusion detection tools. This underscores the complexity of creating reliable and meaningful benchmarks, particularly when dealing with biological material, where variability in sample preparation, sequencing depth, and data quality can substantially affect the outcomes. It was also noted that dataset parameters such as sample type (simulated or real) and read length are important. Most tools give better results for simulated sets due to the characteristics of the set type, lack of noise present in real biological samples. As mentioned earlier, there are inaccuracies in benchmarks related to differences in the truth set. For this reason, such analyses should be approached with a great deal of skepticism. Therefore, a more global comparison at the level of different types of cancers was abandoned.

In this meta-analysis, the history of the Breast cancer cell line dataset was examined, revealing significant variability in its usage across benchmarks. This dataset, published in 2011, includes samples such as BT-474 (SRR064438, SRR064439), SK-BR-3 (SRR064440, SRR064441), KPL-4 (SRR064287), and MCF-7 (SRR064286). Initially, validation indicated the presence of 27 true positive fusions^15^. However, a reanalysis in 2012 revised this number to 40^24^. In 2015, the dataset was used in a benchmark that incorporated 99 true fusions by including fusions from the literature for these cell lines^14^. In 2017, another benchmark utilized the dataset but excluded KPL-4, considering 37 fusions as true positives^11^. In 2018 and 2021, two independent benchmarks^8^,^10^ addressed inconsistencies in the true fusion count by referencing both 27 and 99 fusions simultaneously. In 2019 and 2021, two studies^5^,^9^recognized 53 true fusions, citing validation from other articles. Later in 2021, the dataset was reused with the additional enrichment of MCF-7, and 80 true fusions were taken into account^6^. The inconsistent use of the true fusion count across benchmarks poses a major challenge, as it renders direct comparisons of tool performance difficult or even impossible. This variability underscores the importance of standardizing reference datasets and their annotations to ensure the comparability of results in gene fusion detection studies.

The influence of read length on the quality of the analyses was examined. To ensure a reliable comparison, i.e., having results for at least several tools working on a given dataset, read lengths of 75 bp and 250 bp were excluded in favor of 50 bp and 100 bp. It was observed that, in general, tools tend to perform better with longer read lengths. Given the complications related to real datasets, only simulated datasets were used in this analysis, which eliminates issues related to the biological variability of real samples. It was also noted that read length negatively correlated with performance—shorter read lengths, such as 50 bp, were less effective for properly identifying gene fusions. In some cases, certain tools did not support the analysis of 50 bp samples at all.

The analysis also highlights the importance of tailoring tool selection to specific sample types and experimental conditions. Although the benchmarks, which included a wide range of sequencing protocols, including single-end and paired-end reads, provided a more comprehensive basis for assessing tool performance, too little intersecting data did not allow for a meaningful comparison. Such diversity is essential for researchers who intend to use these tools in a variety of clinical or research settings. Although single-end sequencing is now less popular and its analysis more demanding on tools, sample repositories may still contain a significant number of them (such as the previously mentioned SRA, which contains over 2 million samples or datasets of this type).

A similar situation has befallen the challenge for tools concerning sample size, i.e. the number of reads. Due to the previously mentioned aspects, it cannot be determined whether comparing the results of analyses filtered by dataset size would yield reliable conclusions. It is known that the use of sequencing in the clinic is a constant balance between efficiency and costs, but the appropriate number of reads is important. As mentioned earlier, for WTS these should be higher values, and for targeted sequencing smaller amounts are acceptable. Few benchmarks include datasets from targeted sequencing. Although this is a more popular type of library in clinical sequencing, for the reasons mentioned earlier, there is still no confirmation whether tools for detecting gene fusions behave the same as on full transcriptomes.

From a practical perspective, the results underscore the critical need for standardization in benchmarking practices. For example, while some benchmarks included diverse datasets spanning both simulated and real clinical samples, others did not, limiting their applicability to real-world scenarios. Similarly, inconsistencies in the metrics used for evaluation, such as sensitivity, precision, or F1 score, further complicate cross-comparison of tools. A striking example of the challenges resulting from the lack of standardization is the inconsistent use of the breast cancer cell line dataset across benchmarks. This highlights the importance of not only using well-curated datasets but also ensuring consistency in their interpretation and validation. These challenges reduce reproducibility, make it difficult to reliably compare tools, and limit the applicability of benchmarks to real-world scenarios. Furthermore, computational metrics such as execution time and memory utilization are often overlooked, despite their importance for clinical implementation. Ultimately, this meta-analysis serves as a call to action for the community to establish standardized, comprehensive benchmarks that can help researchers and clinicians select the most appropriate tools for their needs. By addressing the limitations identified in this study, including inconsistencies in dataset utilization and performance metrics, the field can move toward more accurate and reliable gene fusion detection, which is crucial for the advancement of personalized medicine and cancer diagnostics.

To address these limitations and improve the quality of future benchmarks, several steps must be taken. First, establishing standardized gold standards is essential. A universally accepted and biologically verified truth set of gene fusions, combining both simulated and real clinical data, would provide a common reference point for evaluating tool performance. Collaboration among research groups, clinical laboratories, and database maintainers will be crucial in developing and maintaining these standards.

Second, benchmarks should prioritize comprehensive and transparent documentation. This includes providing complete information about tool versions, parameter settings, dataset sources, and preprocessing steps, as well as making all relevant data publicly accessible through repositories. Detailed descriptions of benchmarking workflows should also be included in publications to facilitate reproducibility. Moreover, integrating automated benchmarking pipelines would greatly enhance reproducibility and usability. Open-source pipelines that include tools for dataset preparation, performance evaluation, and result visualization would enable researchers to replicate and extend analyses. Implementing cloud-based or containerized solutions, such as Docker or Singularity, would ensure consistent performance across computational environments.

Aditionally, future benchmarks must include computational metrics alongside traditional accuracy measures. Reporting runtime, memory usage, and scalability will provide a more holistic assessment of tool performance and help identify the best options for large-scale or clinical applications. Additionally, comparative analyses should consider the trade-offs between computational efficiency and detection performance.

Finally, the bioinformatics community should work toward standardizing benchmarking practices. Unified frameworks specifying standardized datasets, evaluation metrics, and reporting formats would reduce variability and improve the comparability of benchmark results. Community-driven efforts, such as consortium-led initiatives or challenges similar to the DREAM Challenges^25^ or FDA-led challenges^26^ in other fields, could foster collaboration and drive innovation in benchmarking practices.

In conclusion, this meta-analysis highlights the variability in gene fusion detection tool performance and underscores the importance of robust benchmark design. Standardizing datasets, improving documentation, and integrating computational metrics into benchmarks will help researchers and clinicians make informed decisions. Addressing these challenges will ultimately improve the accuracy and efficiency of gene fusion detection in clinical and research settings, advancing the field of personalized medicine and cancer diagnostics.

## Methods

### Selected benchmarks

To ensure a comprehensive and unbiased evaluation of gene fusion detection tools, we identified and selected ten publicly available benchmarks from the literature using inclusion criteria described in the section below. Our study compared the performance of 41 tools included in these 10 benchmarks^5–14^. This comprehensive list reflects the wide range of strategies employed for gene fusion detection, emphasizing the necessity of thorough benchmarking to evaluate their performance and suitability for different datasets.

### Inclusion criteria

To identify relevant studies, we performed an extensive search using databases like PubMed and Scopus, employing a variety of keywords, including “gene fusion,” “benchmark,” “genetic tool,” “fusion detection,” and “bioinformatics evaluation.” Only articles published from 2015 onward were considered, and each benchmark had to report performance metrics such as precision, sensitivity, and/or F1 score. Benchmarks that provided additional details on memory usage and time consumption were considered advantageous, but this was not an obligatory criterion. While some benchmarks directly reported precision, sensitivity, and F1 score, others only provided raw data, such as the number of true positives (TP), false positives (FP), and false negatives (FN). In these cases, I derived the required metrics to ensure consistency and comparability across all benchmarks. This approach allowed for a comprehensive and fair evaluation of the performance of gene fusion detection tools. Each benchmark included in this meta-analysis was evaluated based on several criteria, including the number of tools compared (at least 4), dataset type (real or simulated), read length, sample size, and the availability of supporting information such as benchmark code, tool versions, and dataset details.

### Grouping by dataset characteristics

To explore the impact of dataset characteristics on tool performance, we stratified the benchmarks based on:

#### Dataset type

Real (cell culture or patient sample) vs. Simulated Benchmarks were chosen to include both real and simulated datasets. Real datasets were prioritized for their biological relevance and ability to represent genuine gene fusion events. Simulated datasets were included to ensure controlled conditions and evaluate tool performance under specific scenarios.

#### Read length

50 bp, 76 bp, and 100 bp. Short-read sequencing, providing reads of typically 150 bp in length, is supported by a wide range of platforms, dominated by Illumina’s fleet of instruments (NovaSeq, HiSeq, NextSeq, and MiSeq), and is valued for its high accuracy and relative cost-effectiveness. However, sequencing small DNA fragments requires complex algorithms for the reconstruction of longer sequence contigs, potentially resulting in inaccurate and incomplete genome assemblies. This can be further compounded in capture-based sequencing such as exome sequencing where there are inherent limitations in identifying and characterising structural variants^27^

#### Sequencing type

Single-end or Paired-end We selected benchmarks that tested tools across a range of read lengths commonly used in RNA-seq experiments (e.g., 50 bp, 76 bp, 100 bp). Analyzing Sequence Read Archive (SRA) resources for almost 80% of RNA samples “Library Layout” is labeled paired. Although single-end are already losing popularity in the databases on SRA there are still more than 2 million samples sequenced with this parameter.

#### Sample size

Large (*≥*20 million) vs. Small (*<* 20 million) Benchmarks including datasets of varying sizes, from small single-sample datasets to large multi-sample datasets, were considered to evaluate tool scalability. Experiments looking for a more global view of gene expression, and some information on alternative splicing, typically require 30 million to 60 million reads per sample. This range encompasses most published RNA-Seq experiments for mRNA/whole transcriptome sequencing. Targeted RNA expression requires fewer reads. For example, Illumina recommends 3 million reads (Considerations for RNA Seq read length and coverage Illumina^28^)

#### Type of cancer

Relevance to clinical and research applications was included. Benchmarks with datasets derived from clinically relevant cancer types or experimental conditions were considered, as these scenarios represent practical use cases for gene fusion detection tools. Gene fusions play a major role as oncogenic drivers in many cancer types. This insight has immediate consequences for the treatment of patients, because many gene fusions can be addressed therapeutically with targeted drugs. The most prominent examples are fusions between BCR and ABL1 in chronic myeloid leukemia and acute lymphoblastic leukemia, which can be treated effectively using imatinib and related drugs^6^

#### RNA library type

Whole Transcriptome Sequencing (WTS) or targeted sequencing. Currently, more than 10,000 gene fusions have been identified in human cancers. Targeted panels are the most suitable for the clinical practice, require a lower input of starting material, analysis is based on a limited number of clinical valuable targets, are faster and data analysis and result interpretation is easier than with WGS and WTS.^29^

### Methods of analysis

For each tool operating on a specific type of data set, the arithmetic mean and median of F1 score were calculated. We created a series of plots to visually compare the performance of different tools based on F1 score, categorized by dataset type, read length, and sample size. This visualization highlights the variation in tool performance under different conditions. Only tools for which there was sufficient representative data among the benchmark data were included in the plots.

## Author contributions statement

I.O. conceived and conducted the analysis, I.O. and T.G. analyzed the results. T.G. reviewed and edited the manuscript, and provided consultations. All authors reviewed and approved the manuscript.

## Additional information

### Supplementary Materials

Additional data include an Excel sheet with detailed benchmark results, as well as summary tables, such as those listing the tools and datasets used in the analysis.

